# The impact of culture variables on 3D human *in vitro* bone remodeling; a design of experiments approach

**DOI:** 10.1101/2022.11.11.516134

**Authors:** Bregje W.M. de Wildt, Lizzy A.B. Cuypers, Esther E.A. Cramer, Annelieke S. Wentzel, Keita Ito, Sandra Hofmann

## Abstract

Human *in vitro* bone remodeling models, using osteoclast-osteoblast co-cultures, could facilitate the investigation of human healthy (*i*.*e*., balanced) and pathological (*i*.*e*., unbalanced) bone remodeling while reducing the need for animal experiments. Although current *in vitro* osteoclast-osteoblast co-cultures have improved our understanding of bone remodeling, they lack culture method and outcome measurement standardization, which is hampering reproducibility and translatability. Therefore, *in vitro* bone remodeling models could benefit from a thorough evaluation of the impact of culture variables on functional and translatable outcome measures, with the aim to reach ‘healthy’ balanced osteoclast and osteoblast activity. Using a resolution III fractional factorial design, we identified the main effects of commonly used culture variables on bone turnover markers in a robust *in vitro* human bone remodeling model. Our model was able to capture physiological quantitative resorption – formation coupling along all conditions, whereby remodeling could be enhanced by external stimuli. Especially culture conditions of two runs showed promising results, where conditions of one run could be used as a high bone turnover system and conditions of another run as a self-regulating system as the addition of osteoclastic and osteogenic differentiation factors was not required for remodeling. The results generated with our *in vitro* model allow for better translation between *in vitro* studies and towards *in vivo* studies, for improved preclinical bone remodeling drug development.

**Graphical abstract:** 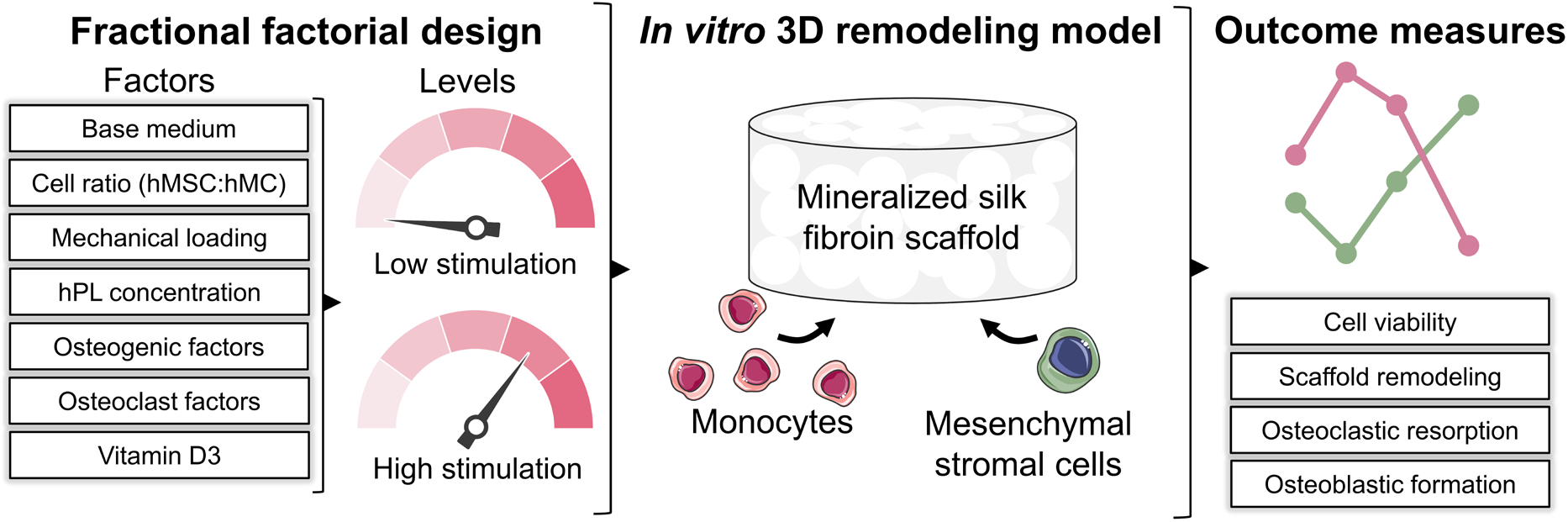

## 1. Introduction

Bone is a highly dynamic tissue continuously remodeled by bone resorbing osteoclasts, bone forming osteoblasts, and regulating osteocytes. Physiological or healthy bone remodeling requires balanced formation and resorption. A shift in this balance, towards more resorption or formation, is a hallmark for bone pathologies like osteoporosis or osteopetrosis, respectively. Studies of these bone pathologies and their treatment testing are routinely performed in animal models. These animal models often represent human physiology insufficiently, which is likely one of the reasons that only 8-10% of preclinically developed drugs are approved for regular clinical use [1–3]. Human *in vitro* bone remodeling models could facilitate the investigation of human healthy and pathological bone remodeling while addressing the principle of reduction, refinement, and replacement of animal experiments (3Rs) [4,5].

A co-culture of osteoclasts and osteoblasts is minimally needed to mimic the bone remodeling process *in vitro* [6]. For these co-cultures, human monocytes (hMCs) and mesenchymal stromal cells (hMSCs) are most frequently used as progenitor cells which are in culture differentiated into osteoclasts and osteoblasts (and eventually osteocytes), respectively [6]. To stimulate hMCs and hMSCs to undergo differentiation and subsequently study *in vitro* remodeling, a variety of culture conditions and outcome measures are used which differ for each research group and/or study aim [6,7]. Variations in culture protocols include *e*.*g*., different cell ratios, different base media, the use of osteogenic/osteoclast supplements and their respective concentrations, and the application of mechanical load [7]. These culture variables could lead to unequal stimulation of osteoblasts and osteoclasts which might cause an unhealthy resorption-formation balance. Moreover, outcome measures of current models often include only the evaluation of osteoclast and/or osteoblast markers with gene expression analysis or enzymatic activity assays rather than their functionality to resorb and form a bone-like extracellular matrix *in vitro* [7]. However, these cell-matrix interactions are often the main outcome measure for *in vivo* studies (*i*.*e*., the evaluation of bone structure change using X-ray based methods or the measurement of blood collagen formation and degradation markers). Thus, current *in vitro* osteoclast-osteoblast co-cultures have improved our understanding of bone remodeling, they however lack culture method and outcome measurement standardization, hampering reproducibility and translatability to *in vivo* animal models and *in vivo* human data. In this regard, *in vitro* bone remodeling models could benefit from a thorough evaluation of the impact of culture variables on functional and translatable outcome measures, with the aim to reach ‘healthy’ balanced osteoclast and osteoblast activity.

Researchers have already attempted to study the influences of culture variables on human osteoblast-osteoclast co-cultures. For example, studies looked at the influence of cell ratio on osteoclast formation [8], osteogenic factor addition and timing on osteogenic and osteoclastic differentiation [9,10], and the replacement of the culture supplement fetal bovine serum (FBS) by serum free medium [11], or human platelet lysate (hPL) [12] on osteoclastic resorption. As such, most studies analyze the influence of only one culture variable on *in vitro* remodeling outcomes while a specific combination of multiple variables might lead to improved results. A fractional factorial design of experiments (DoE) approach could facilitate the time-efficient evaluation of the impact of multiple culture variables on *in vitro* remodeling outcomes [13]. A fractional factorial design assumes that higher order interaction effects are insignificant. Factor main effects can therefore be evaluated using a fraction of the required experiments of a full factorial design study. While regularly used in most engineering fields, the DoE approach is not often employed for bioengineering. For bioengineering, DoE have been employed for *e*.*g*., the optimization of biomaterials [14,15], or the optimization of culture conditions for improved human pluripotent stem cell expansion [16], osteogenic differentiation of adipose derived hMSCs [17], or vascular network formation in bone-like constructs [18]. In this study, we used a fractional factorial design to evaluate the impact of culture variables on functional and translatable outcome measures in an *in vitro* remodeling model (Figure 1) [19]. The influence of commonly used culture variables [7], including base medium, cell ratio, mechanical loading, hPL concentration, osteogenic differentiation factors, osteoclast differentiation factors and 1,25-dihydroxyvitamin D3, on mainly non-destructive bone remodeling outcomes was evaluated over a period of 28 days. Outcome measures included sequential (registered) micro-computed tomography (μCT) images and the longitudinal evaluation of resorption by tartrate-resistant acid phosphatase (TRAP) and cathepsin K quantification, and formation by alkaline phosphatase (ALP) and pro-collagen 1 c-terminal propeptide (PICP) quantification as commonly used bone turnover biomarkers [20]. Besides, cell metabolic activity and cell death were longitudinally monitored. With this study, we aimed at finding culture conditions that equally support osteoclastic and osteogenic differentiation of hMCs and hMCSs, respectively, followed by balanced *in vitro* remodeling.

**Figure 1.**
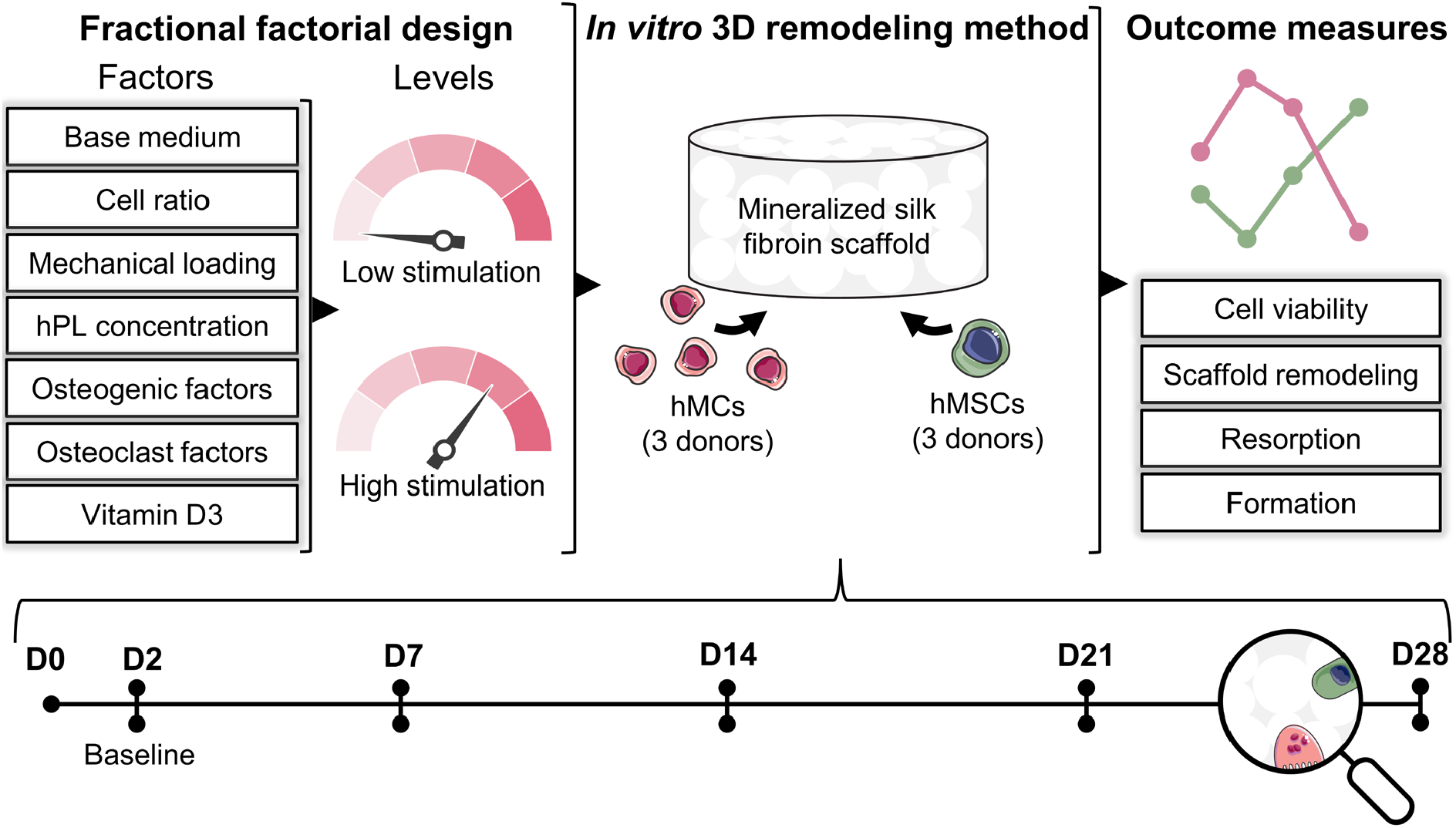
Experimental set-up of the study. A fractional factorial design was used to study the influences of the culture variables (base medium, cell ratio, mechanical loading, hPL concentration, osteogenic differentiation factors, osteoclastic differentiation factors and 1,25-dihydroxyvitamin D3) on cell viability, scaffold remodeling balance, osteoclastic resorption and osteoblastic formation in an *in vitro* bone remodeling model. A co-culture of hMCs and hMSCs was maintained for a period of 28 days during which remodeling was tracked non-destructively. Abbreviations: human platelet lysate (hPL), human monocytes (hMCs), human mesenchymal stromal cells (hMSCs), day (D). Parts of the figure were modified from Biorender.com (https://biorender.com/, accessed on 6 October 2022) and Servier Medical Art, licensed under a Creative Common Attribution 3.0 Generic License (http://smart.servier.com/, accessed on 8 July 2021).

## 2. Materials and Methods

### 2.1 Factor selection and experimental matrix creation

To select the parameters to be tested in the DoE set-up, our database, as part of a systematic review, with culture conditions of all identified *in vitro* bone remodeling models was consulted [7]. From this database, the following culture variables were identified: culture substrate/material, cell type, seeding density, base medium, co-culture cell ratio, biomechanical environment, serum supplement, osteogenic differentiation factors (*i*.*e*., dexamethasone, *β*-glycerophosphate and ascorbic acid), osteoclast differentiation factors (*i*.*e*., receptor activator of nuclear factor κB ligand (RANKL) and macrophage colony-stimulating factor (M-CSF)), and the use of additional factors of which 1,25-dihydroxyvitamin D3 was most commonly used. For this study, the influence of culture substrate/material, cell type and the hMSC seeding density were not included as these factors form the base of our *in vitro* model that was used to study the effect of the other factors [19]. For the other factors, two levels (*i*.*e*., low stimulation and high stimulation) were assigned (Table 1). For base medium, levels were *α*-MEM and DMEM as most commonly used co-culture base media [7]. For cell ratio, ratios of 1:2 and 1:5 (hMSCs:hMCs) were included. For mechanical loading, static and dynamic loading were included, using a custom made spinner flask bioreactor at 300 RPM to apply fluid shear stress [21]. As a serum supplement, hPL was used at 5% and 10% concentration since the most commonly used FBS can inhibit osteoclast resorption [11,12]. For osteogenic supplements, ascorbic acid was used in all conditions as a requirement for collagen synthesis. Dexamethasone was added in a low concentration of 10 nM, which is a commonly used concentration for co-cultures and believed to be the physiological glucocorticoid concentration known to stimulate osteogenesis and osteoclastogenesis [22,23]. The other commonly used co-culture dexamethasone concentration of 100 nM was used for high stimulation [7]. *β*-glycerophosphate was only added in the high stimulation condition at the most commonly used concentration of 10 mM [7]. As the material used in this study contained hydroxyapatite, resorption was expected to release sufficient phosphate for osteogenic differentiation and mineralization to leave out *β*-glycerophosphate in the low stimulation condition. M-CSF and RANKL were only added in high stimulation conditions at commonly used concentrations of 50 ng/ml [7]. The concentration of 1,25-dihydroxyvitamin D3 was set at the common concentration 10 nM in high stimulation conditions, whereas in low stimulation conditions no 1,25-dihydroxyvitamin D3 was added (Table 1).

**Table 1.** Evaluated factors and their corresponding levels.

|  | Factor |  | Level |  | Unit |
| --- | --- | --- | --- | --- | --- |
|  |  |  | Low | High |  |
| **A** | Base medium | | α-MEM | DMEM | - |
| **B** | Cell ratio |  | 1:2 | 1:5 | - |
| **C** | Mechanical loading* |  | 0 | 300 | RPM |
| **D** | hPL concentration |  | 5 | 10 | % |
| **E** | Osteogenic factors** | Dexamethasone | 10 | 100 | nM |
| | | β-glycerophosphate | 0 | 10 | mM |
| **F** | Osteoclast factors | M-CSF | 0 | 50 | ng/ml |
|  |  | RANKL*** | 0 | 50 | ng/ml |
| **G** | 1,25-dihydroxyvitamin D3 |  | 0 | 10 | nM |

Abbreviations: human platelet lysate (hPL), macrophage colony-stimulating factor (M-CSF), receptor activator of nuclear factor κB ligand (RANKL).
\*applied from day 2 in culture
\*\*ascorbic acid was present in all cultures
\*\*\*added from day 2 in culture

For the resulting 7 factors with 2 levels, a resolution III fractional factorial design was randomly created using R (version 4.1.2) [24] with the Rcmdr DoE plugin (version 0.12-3, Ulrike Groemping) [25], leading to 8 experimental runs (Table 2). Resolution ≥ III designs are considered appropriate for screening purposes. An additional run was included in which all factors had level low, which served as a negative control (run 9), a positive control (all factors level high) was already part of the design (run 7) (Table 2).

**Table 2.** Experimental matrix

| Run | A | B | C | D | E | F | G |
| --- | --- | --- | --- | --- | --- | --- | --- |
| 1 | α-MEM | 1:5 | Dynamic | 5% hPL | | OCL fact | |
| 2 | α-MEM | 1:5 | Static | 5% hPL | OG fact | | vitD3 |
| 3 | DMEM | 1:5 | Static | 10% hPL |  |  |  |
| 4 | α-MEM | 1:2 | Dynamic | 10% hPL | | | vitD3 |
| 5 | DMEM | 1:2 | Static | 5% hPL |  | OCL fact | vitD3 |
| 6 | DMEM | 1:2 | Dynamic | 5% hPL | OG fact |  |  |
| 7 | DMEM | 1:5 | Dynamic | 10% hPL | OG fact | OCL fact | vitD3 |
| 8 | α-MEM | 1:2 | Static | 10% hPL | OG fact | OCL fact | |
| 9 | α-MEM | 1:2 | Static | 5% hPL | | | |

Abbreviations: human platelet lysate (hPL), osteogenic differentiation factors (OG fact), osteoclast differentiation factors (OCL fact), 1,25-dihydroxyvitamin D3 (vitD3). A = base medium, B = cell ratio, C = mechanical loading, D = human platelet lysate concentration, E = osteogenic factors, F = osteoclast factors, G = 1,25-dihydroxyvitamin D3

### 2.2 Scaffold fabrication

Bombyx mori L. silkworm cocoons were degummed by boiling them in 0.2 M Na_2_CO_3_ (S-7795, Sigma-Aldrich, Zwijndrecht, The Netherlands) for 1 h. Air-dried silk fibroin (SF) was dissolved in 9 M LiBr (199870025, Acros, Thermo Fisher Scientific, Breda, The Netherlands), filtered, and dialyzed against ultra-pure water (UPW) for 36 h using SnakeSkin Dialysis Tubing (molecular weight cut-off: 3.5 K, 11532541, Thermo Fisher Scientific). The dialyzed SF solution was frozen at -80 ºC and lyophilized for 7 days. Lyophilized SF was dissolved in hexafluoro-2-propanol (003409, Fluorochem, Hadfield, UK) at a concentration of 17% (w/v) and casted in scaffold molds containing NaCl granules with a size of 250-300 μm as template for the pores. Molds were covered to improve the SF blending with the granules. After 3 h, covers were removed from molds, and hexafluoro-2-propanol was allowed to evaporate for 7 days whereafter *β-*sheets were induced by submerging SF-salt blocks in 90% MeOH for 30 min. SF-salt blocks were cut into discs of 3 mm height with a Accutom-5 (04946133, Struer, Cleveland, OH, USA). NaCl was dissolved for 48 h from the scaffolds in UPW, resulting in porous sponges. From these sponges, scaffolds were punched with a 5 mm diameter biopsy punch.

### 2.3 Scaffold mineralization

SF scaffolds were mineralized as previous described [19], using a mineralization solution with 10x SBF [26] and 100 μg/ml poly-aspartic acid (pAsp, P3418, Sigma-Aldrich). Briefly, a 10x SBF stock was prepared. Just prior to mineralization, mineralization solution was prepared by adding 100 μg/ml pAsp to 10x SBF, followed by the addition of NaHCO_3_ until a final concentration of 10 mM, both under vigorous steering. Scaffolds were incubated with 8.6 ml mineralization solution for 2 weeks at 37 ºC on an orbital shaker at 150 RPM in mineralization solution with a solution replenishment after 1 week. After mineralization, scaffolds were washed 3 × 15 min in an excess of UPW. Scaffolds were sterilized by autoclaving in phosphate buffered saline (PBS) at 121 ºC for 20 min.

### 2.4 Cell culture experiments

#### 2.4.1 Monocyte isolation

Peripheral blood mononuclear cells (PBMCs) were isolated from human peripheral blood buffy coats (Sanquin, Eindhoven, The Netherlands; collected under their institutional guidelines and with informed consent per Declaration of Helsinki) of three healthy donors. The buffy coats (∼50 ml each) were diluted with 0.6% w/v sodium citrate in PBS (citrate-PBS) until a final volume of 200 ml and layered per 25 ml on top of 10 ml Lymphoprep^TM^ (07851, StemCell technologies, Köln, Germany) in 50 ml centrifugal tubes. After density gradient centrifugation (20 min at 800x g, lowest break), PBMCs were collected, resuspended in citrate-PBS, and washed four times in citrate-PBS supplemented with 0.01% bovine serum albumin (BSA, 10735086001, Merck KGaA, Darmstadt, Germany). PBMCs were frozen at 10^5^ cells/ml in freezing medium containing RPMI-1640 (RPMI, A10491, Thermo Fisher Scientific), 20% fetal bovine serum (FBS, BCBV7611, Sigma-Aldrich) and 10% dimethyl sulfoxide (DMSO, 1.02952.1000, VWR, Radnor, PA, USA) and stored in liquid nitrogen until further use. Before hMC isolation, PBMCs were thawed, collected in hMC isolation medium containing RPMI, 10% FBS (BCBV7611, Sigma-Aldrich) and 1% penicillin-streptomycin (p/s, 15070063, Thermo Fisher Scientific), and after centrifugation resuspended in isolation buffer (0.5% w/v BSA in 2mM EDTA-PBS). hMCs were enriched from PBMCs with manual magnetic activated cell separation (MACS) using the Pan Monocyte Isolation Kit (130-096-537, Miltenyi Biotec, Leiden, Netherlands) and LS columns (130-042-401, Miltenyi Biotec) according to the manufacturer’s protocol, and directly used for experiments.

#### 2.4.2 hMSC isolation, expansion and seeding

hMSCs were isolated from human bone marrow of three healthy donors (1M-125, Lonza, Walkersville, MD, USA, collected under their institutional guidelines and with informed consent) and characterized for surface markers and multilineage differentiation, as previously described [27]. Bone-marrow derived hMSCs (hBMSCs) were frozen at passage 3 or 4 with 5*10^6^ cells/ml in freezing medium containing FBS (BCBV7611, Sigma-Aldrich) with 10% DMSO and stored in liquid nitrogen until further use. Before experiments, hBMSCs were thawed, collected in high glucose DMEM (hg-DMEM, 41966, Thermo Fisher Scientific), seeded at a density of 2.5*10^3^ cells/cm^2^ and expanded in expansion medium containing hg-DMEM, 10% FBS (BCBV7611, Sigma-Aldrich), 1% Antibiotic Antimycotic (anti-anti, 15240, Thermo Fisher Scientific), 1% Non-Essential Amino Acids (11140, Thermo Fisher Scientific), and 1 ng/ml basic fibroblast growth factor (bFGF, 100-18B, PeproTech, London, UK) at 37 ºC and 5% CO_2_. After 7-10 days, at around 80% confluence, cells were detached using 0.25% trypsin-EDTA (25200, Thermo Fisher Scientific) and seeded onto scaffolds at passage 4 or 5.

#### 2.4.3 hMC-hBMSC co-culture on mineralized SF scaffolds

hBMSCs were seeded at a density of 0.5*10^6^ cells per scaffold and seeding was performed dynamically [28] in 50 ml tubes on an orbital shaker at 150 RPM in expansion medium. After 6 hours, scaffolds were transferred to 24-wells plates and hMCs were seeded in hMC isolation medium at a density of 1*10^6^ or 2.5*10^6^ cells/20 μl (dependent on the experimental run) by pipetting 20 μl of cell suspension onto the scaffolds. Cells were allowed to attach for 90 min at 37 ºC and every 20 minutes a small droplet of medium from the respective experimental run was added. Per experimental run, 4 different hMC and hBMSC donor combinations (1 – 3 repeats per donor combination) were seeded on *N* = 8 scaffolds. The cell-loaded scaffolds were subsequently placed in custom-made spinner flask bioreactors (2 donor combinations on 4 scaffolds per bioreactor, 2 bioreactors per experimental run) and cultured statically or dynamically for 28 days at 37 ºC and 5% CO_2_ in their respective medium (Table S1 and S2). Medium was replaced 3x per week and medium samples were collected on day 2 and weekly from day 7, and stored at -80 ºC. Constructs were sacrificed for analyses after 28 days of culture.

### 2.5 Analyses

#### 2.5.1 μCT

On day 2, 7, 14, 21 and 28, scaffolds (*N* = 8 per run) were scanned and later analyzed with a *μ*CT100 imaging system (Scanco Medical, Brüttisellen, Switzerland). Scanning was performed with an energy level of 45 kVp, intensity of 200 μA, integration time of 300 ms and with twofold frame averaging. To reduce part of the noise, a constrained Gaussian filter was applied to all scans with filter support 1 and filter width sigma 0.8 voxel. Follow-up images were registered to the image of the previous time point, such that voxels at the surface of the scaffold were categorized into resorption site, formation site, or unchanged/quiescent site [29]. The scaffold was segmented at a global threshold of 24% of the maximum grayscale value and remodeled scaffold surface was segmented at a global threshold of 7.5% of the maximum grayscale value. This threshold was chosen after registration of cell-free construct images in such a way that resorption and formation were below ∼1.5% of the total volume to ensure that remodeled volume was caused by the cells instead of by noise. To further reduce noise, only a minimum cluster of 2 resorbed or formed voxels were included in the analyses. For illustration purposes, day 28 images were also registered to day 2 images for the total resorption and formation visualization.

#### 2.5.2 PrestoBlue^TM^ assay

On day 2, 7, 14, 21 and 28, scaffolds were incubated with 10% v/v PrestoBlue^TM^ (A13262, Thermo Fisher Scientific) in their respective medium (without supplementation of 1,25-dihydroxyvitamin D3, osteogenic or osteoclast factors) within their bioreactors for 25 min at 37 ºC in the dark. Samples (*N* = 8, 4 technical repeats from 2 bioreactors with each 4 scaffolds per run) were pipetted in duplo in black 96-wells assay plates. Fluorescence (excitation: 530/25 nm, emission 590/35 nm) was measured with a plate reader (Synergy^TM^ HTX, Biotek). Measured fluorescence was corrected for blank medium samples.

#### 2.5.3 Lactate dehydrogenase activity (LDH)

On cell supernatants from day 2, 7, 14, 21 and 28, LDH activity was measured (*N* = 8, 4 technical repeats from 2 bioreactors with each 4 scaffolds per run). A 100 μl supernatant sample or NADH (10107735001, Sigma-Aldrich) standard was in duplo incubated with 100 μl LDH reaction mixture (11644793001, Sigma-Aldrich) in 96-wells assay plates. Absorbance was measured after 5, 10 and 20 min at 490 nm using a plate reader, and LDH activity was calculated between the 10 and 20 min reactions, using standard curve absorbance values.

#### 2.5.4 TRAP activity

On cell supernatants from day 2, 7, 14, 21 and 28, TRAP activity was quantified (*N* = 8, 4 technical repeats from 2 bioreactors with each 4 scaffolds per run). A 10 μl supernatant sample or p-nitrophenol standard was in duplicate incubated with 90 μl p-nitrophenyl phosphate buffer (1 mg/ml p-nitrophenyl phosphate disodium hexahydrate (71768, Sigma-Aldrich), 0.1 M sodium acetate, 0.1% Triton X-100 and 30 μl/ml tartrate solution (3873, Sigma-Aldrich) in PBS) in 96-wells assay plates for 90 min at 37 ºC. To stop the reaction, 100 μl 0.3 M NaOH was added. Absorbance was read at 405 nm using a plate reader and absorbance values were converted to TRAP activity (converted p-nitrophenyl phosphate in nmol/ml/min) using standard curve absorbance values.

#### 2.5.5 Cathepsin K activity

On cell supernatants from day 2, 7, 14, 21 and 28, Cathepsin K activity was quantified (*N* = 8, 4 technical repeats from 2 bioreactors with each 4 scaffolds per run). A 50 μl supernatant sample or aminomethylcoumarin (A9891, Sigma-Aldrich) standard was in duplo incubated with 50 μl substrate working solution (100 μM Z-LR-AMC (BML-P229-0010, Enzo Life Sciences, Bruxelles, Belgium), 0.1 M sodium acetate trihydrate, 4 mM EDTA and 4 mM DTT at pH 5.5 in UPW) in 96-wells assay plates for 30 min at 37 ºC. Fluorescence (excitation: 360/40 nm, emission 460/40 nm) was measured with a plate reader and values were converted to Cathepsin K activity (converted Z-LR-AMC in μmol/ml/min) using standard curve fluorescence values.

#### 2.5.6 PICP quantification

On cell supernatants from day 2, 7, 14, 21 and 28, PICP as collagen formation product was quantified using an enzyme-linked immunosorbent assay (ELISA, MBS2502579, MyBioSource, San Diego, CA, USA) according to the manufacturer’s protocol. Samples (*N* = 2, one sample per bioreactor with each 4 scaffolds per run) were added in triplicate to anti-human PICP coated microwells. After 90 min incubation at 37 ºC, samples were replaced by biotinylated antibody solution followed by 60 min incubation at 37 ºC. After thorough washing, HRP-conjugate solution was added, and plates were incubated for 30 min at 37 ºC. Wells were again washed, and substrate reagent was added followed by 15 min incubation in the dark at 37 ºC. To stop the reaction, stop solution was added and absorbance was measured at 450 nm in a plate reader. Absorbance values were converted to PICP concentrations using standard curve absorbance values.

#### 2.5.7 Alkaline phosphatase activity

On day 28, scaffolds (*N* = 4 per run) were washed in PBS and disintegrated using 2 steel balls and a mini-beadbeater^TM^ (Biospec, Bartlesville, OK, USA) in 500 μl cell lysis buffer containing 0.2% (v/v) Triton X-100 and 5 mM MgCl_2_. ALP activity in cell lysates was determined by adding 20 μl of 0.75 M 2-amino-2-methyl-1-propanol (A65182, Sigma-Aldrich) to 80 μl sample in 96-wells assay plates. Subsequently, 100 μl substrate solution (10 mM p-nitrophenyl-phosphate (71768, Sigma-Aldrich) in 0.75 M 2-amino-2-methyl-1-propanol) was added and wells were incubated at room temperature for 15 minutes. To stop the reaction, 100 μl 0.2 M NaOH was added. Absorbance was measured with a plate reader at 450 nm and these values were converted to ALP activity (converted p-nitrophenyl phosphate in μmol/ml/min) using standard curve absorbance values.

#### 2.5.8 Scanning electron microscopy (SEM)

On day 28, scaffolds (*N* = 1-3 per run) were fixed in 2.5% glutaraldehyde in 0.1 M sodium cacodylate buffer (CB) for 4 h and then washed in CB. Samples were dehydrated with graded ethanol series (37%, 67%, 96%, 3 × 100%, 15 minutes each), followed by a hexamethyldisilazane (HDMS)/ethanol series (1:2, 1:1, 2:1, 3 × 100% HDMS, 15 minutes each). Samples were coated with 20 nm gold and imaging was performed in high vacuum, at 10 mm working distance, with a 5kV electron beam (Quanta 600F, FEI, Eindhoven, The Netherlands).

#### 2.5.9 Histochemical analysis and confocal microscopy

On day 28, scaffolds (*N* = 1-3 per run) were fixed overnight in 3.7% neutral buffered formaldehyde, washed in PBS, permeabilized for 30 min in 0.5% Triton X-100 in PBS and stained overnight with 1 μmol/mL CNA35-OG488 [30] and 0.2 nmol/ml OsteoSense^TM^ 680 (NEV10020EX, PerkinElmer, Waltham, MA, USA) at 4 ºC to visualize collagen and hydroxyapatite, respectively. After washing with PBS, samples were incubated for 1 h with 1 μg/ml DAPI and 50 pmol Atto 550-conjugated Phalloidin (19083, Sigma-Aldrich). Samples were washed and imaged in PBS images were acquired with a confocal laser scanning microscope (Leica TCS SP8X, 40x/0.95 HC PL APO objective).

### 2.6 Statistical analyses

Statistical analyses were performed, and graphs were prepared in GraphPad Prism (version 9.3.0, GraphPad, La Jolla, CA, USA) and R (version 4.1.2) [24] with the Rcmdr DoE plugin (version 0.12-3, Ulrike Groemping) [25]. Statistical analyses were only done for day 21 data, as osteoclasts have a limited lifespan of about 14-21 days [31,32], and osteogenesis takes about 14-21 days [33]. ALP data was analyzed at day 28, as endpoint analysis. For the comparison between different experimental runs, data were tested for normality in distributions and equal variances using Shapiro-Wilk tests and Levene’s tests, respectively. When these assumptions were met, mean ± standard deviation are presented and a one-way ANOVA was performed followed by Holm-Šídák’s post hoc tests with adjusted *p*-values for multiple comparisons, in which experimental runs were pairwise compared with the negative (run 9) and positive (run 7) control. Other data are presented as median ± interquartile range and were tested for differences with the non-parametric Kruskal-Wallis test with Dunn’s post hoc tests with adjusted p-value for pairwise comparisons with the positive and negative control. To quantify the resorption-formation coupling, a spearman correlation coefficient was calculated for the μCT outcomes resorbed mineralized volume – formed mineralized volume and for the supernatant outcomes TRAP activity – PICP concentration. As part of the fractional factorial design analysis, factor main effect plots and effect normal plots were prepared for effect visualization and factor significance [25], respectively. Due to failed μCT registration, some experimental runs missed 1 – 2 out of 8 samples for mineral formation and resorption quantification. To allow for a balanced factorial design analysis, the average of the respective experimental runs was included as additional 1 – 2 sample values. A *p*-value of <0.05 was considered statistically significant.

## 3. Results

### 3.1 Cell viability

From day 7 to day 28, metabolic activity increased for all experimental runs (Figure 2A). When comparing the metabolic activity on day 21 of each run with the metabolic activity of the positive (only high stimulation) and negative (only low stimulation) control, a statistically significant lower metabolic activity was found in the positive control (run 7) when compared with run 1, 2, 3 and the negative control (run 9) (Figure 2B). The negative control also had a statistically significant higher metabolic activity than run 4 (Figure 2B). LDH activity in the supernatant, as a measure for cell death, initially decreased for most experimental runs while towards day 21 it tended to stabilize or increase slightly (Figure 2C). While the metabolic activity was relatively low in the positive control, a statistically significant higher day 21 LDH activity was found when comparing the positive control with run 1, 4, 5 and the negative control (run 9) (Figure 2D). Overall, cell death was lowest in the negative control with a statistically significant difference when comparing to run 3 and 8 (Figure 2D). Factor main effect plots indicated the influence of the culture variables and their corresponding levels on metabolic activity and cell death (Figure 2E). No significant contribution to metabolic activity nor cell death was found from one of the factors. From the metabolic activity main effect plots and normal effect plots, a high cell ratio (1:5) tended to positively influence metabolic activity (Figure 2E and S1). This is likely because of the higher total number of cells as the hBMSC seeding density remained constant. Interestingly, all other factors seemed to negatively impact metabolic activity when high stimulation was applied (Figure 2E and S1). A high concentration of hPL tended to increase LDH activity (Figure 2E and S1), likely caused by the presence of LDH in human platelets [34].

**Figure 2.**
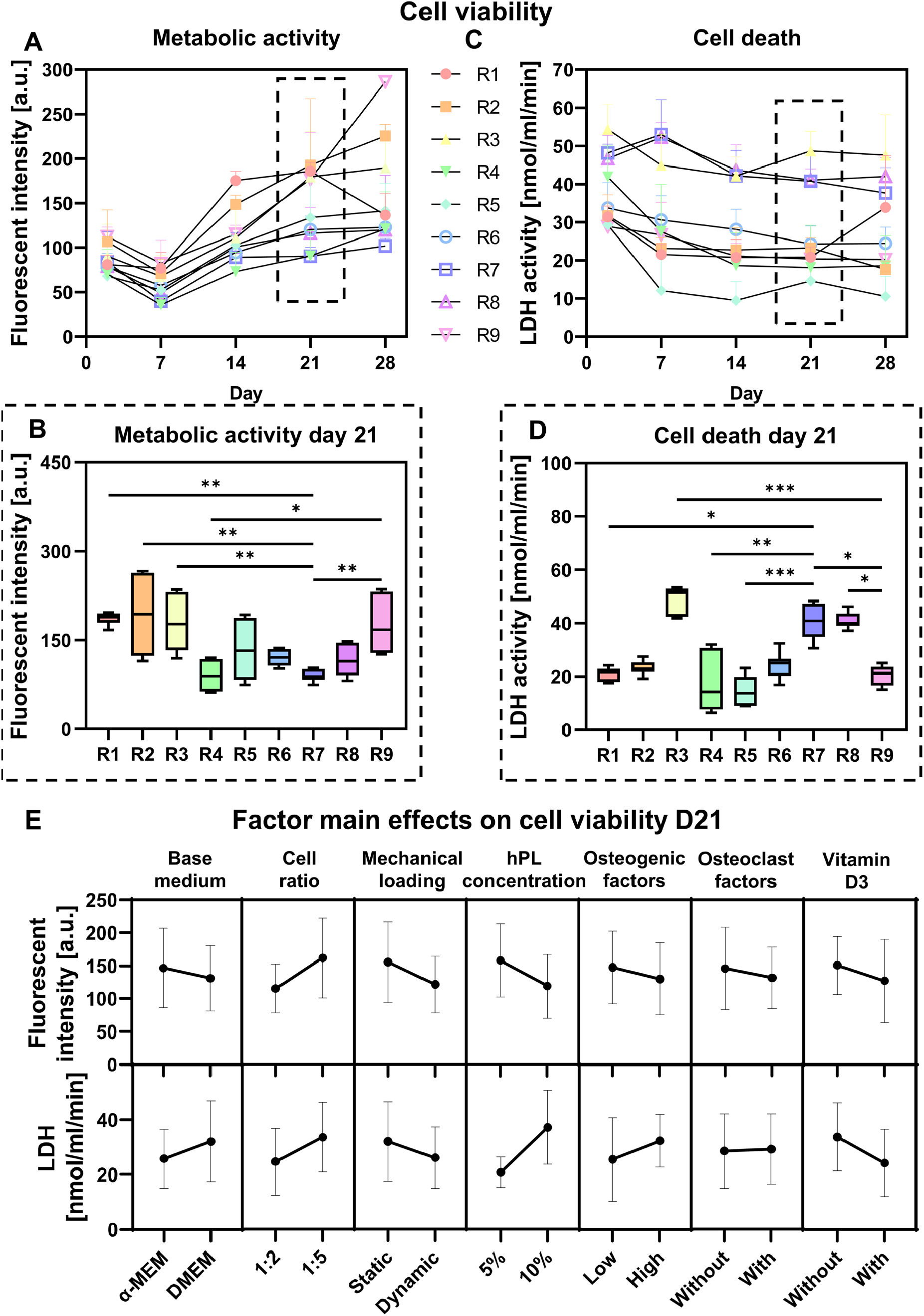
Cell viability testing of experimental runs. (**A**) Metabolic activity measurements using PrestoBlue^TM^ on day 2, 7, 14, 21, and 28. (**B**) Day 21 metabolic activity measurements, *p*<0.05 (Kruskal-Wallis and Dunn’s post hoc tests). (**C**) Cell death measured by LDH release in the medium on day 2, 7, 14, 21, and 28. (**D**) Day 21 cell death measurements, *p*<0.05 (Kruskal-Wallis and Dunn’s post hoc tests). (**E**) Factor main effects and standard deviations on day 21 cell viability outcome measures. (\**p*<0.05, \*\**p*<0.01, \*\*\**p*<0.001). Abbreviations: lactate dehydrogenase activity (LDH), run (R), human platelet lysate (hPL).

### 3.2 Scaffold remodeling

When visualizing mineral resorption and formation sites after registration of day 28 μCT images to day 2 images, remodeling was observed in all experimental runs (Figure 3A). While in the negative control (run 9) limited remodeling was observed, the positive control (run 7) showed extensive remodeling with mostly mineral formation (Figure 3A). Quantification of the percentage formed, resorbed and quiescent (unremodeled) mineral between day 2 -7, 7 - 14, 14 - 21, and 21 - 28, allowed for calculating the balance between formed and resorbed mineral (*i*.*e*., mineral formation – mineral resorption) (Figure 3B). Remarkably, in all experimental runs more formation than resorption was observed for most time points (Figure 3B). Over time, the negative control (run 9) showed most balanced remodeling with limited net resorption or formation, while in the positive control (run 7) a relatively high net formation was observed at all time points (Figure 3B). From day 14 - 21, there was indeed a significantly higher net formation in the positive control when compared to run 3, 5, 8, and the negative control (Figure 3C), which have in common that they were all cultured statically. In addition to the positive control, run 6 also had a significantly higher net formation than the negative control (Figure 3C). Comparison of the quiescent scaffold mineralized volume revealed that most scaffold was remodeled in the initial 21 days, observed by a small increase in quiescent volume after day 21 consistent for all experimental runs (Figure 3D). Overall, most scaffold remodeling seemed to take place in the positive control and least remodeling in the negative control (Figure 3D). From day 14 – 21, least quiescent scaffold mineralized volume was observed in run 1, with statistically significant less quiescent volume than the negative control (run 9) (Figure 3E).

**Figure 3.**
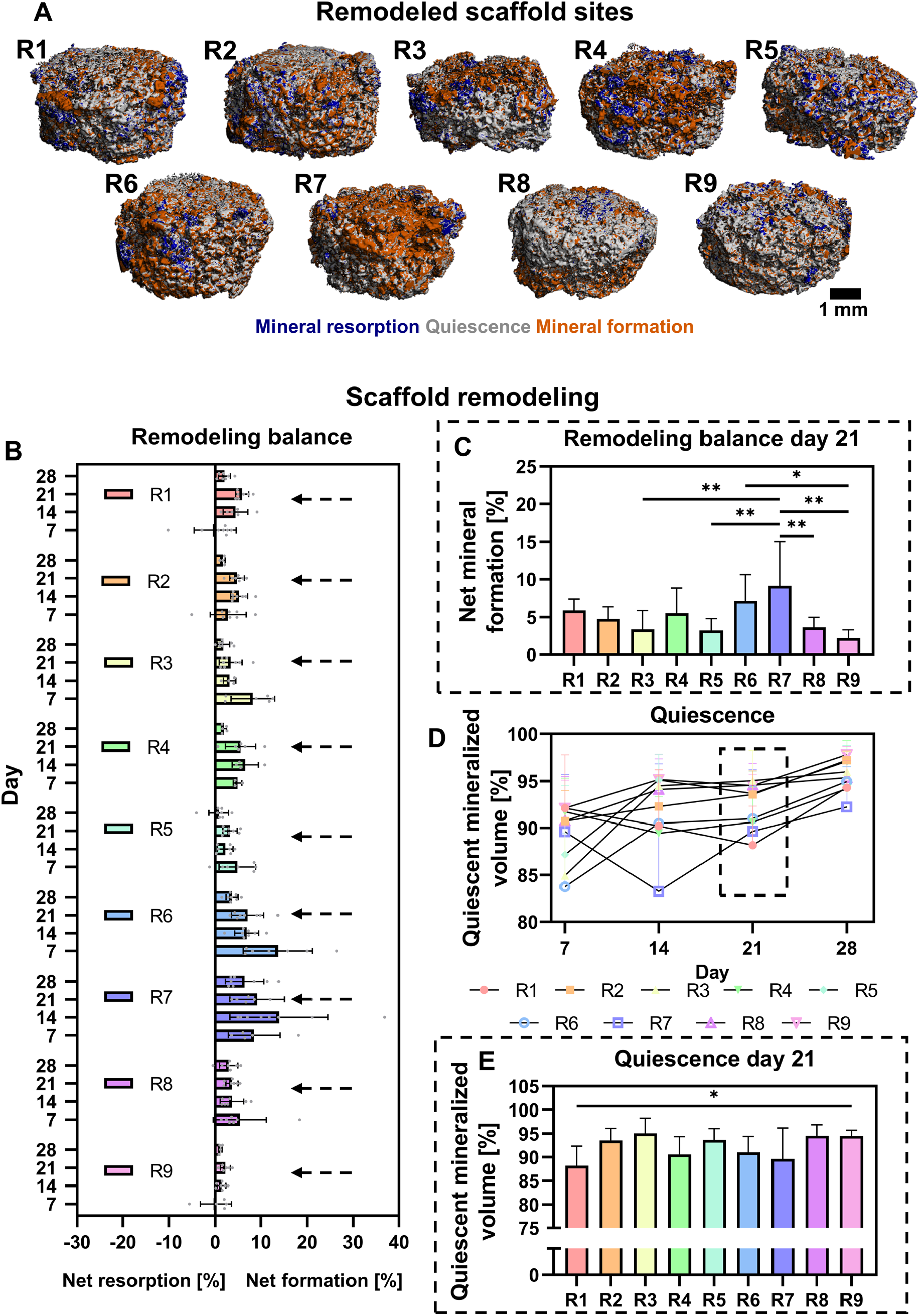
Scaffold remodeling of experimental runs. (**A**) Remodeled scaffolds sites between day 2 and 28, obtained with μCT. (**B**) μCT based formed mineral – resorbed mineral as measure for remodeling balance of day 2-7, 7-14, 14-21, and 21-28. (**C**) Day 14-21 remodeling balance, *p*<0.05 (One-way ANOVA and Holm-Šídák’s post hoc tests). (**D**) μCT based quiescent mineral or unremodeled scaffold mineral of day 2-7, 7-14, 14-21, and 21-28. (**E**) Day 14-21 quiescent mineral, *p*<0.05 (One-way ANOVA and Holm-Šídák’s post hoc tests). (\**p*<0.05, \*\**p*<0.01). Abbreviations: run (R).

### 3.3 Osteoclastic resorption

Over the culture period, experimental run 1 showed consistently most resorbed mineralized volume (Figure 4A). In both the positive (run 7) and negative (run 9) controls, limited resorption was observed (Figure 4A). When comparing the resorbed mineralized volume from day 14 - 21, statistically significant more mineral resorption was observed in experimental run 1 when compared with the positive control (Figure 4B). Interestingly, although not significant, the osteoclast factor-lacking run 4 also showed a relatively high resorbed mineralized volume. For run 1 and 4, the high resorbed mineralized volume was reflected in a relatively high TRAP activity and for run 1 also Cathepsin K activity (Figure 4C+E). Although limited resorbed mineralized volume was observed for the positive control (run 7), a relatively high TRAP activity was measured (Figure 4C). On day 21, when compared to the negative control (run 9), a statistically significant higher TRAP activity was measured in run 1 and 7 (Figure 4D). When compared to the positive control (run 7), a statistically significant lower TRAP activity was found in run 2 and 5 (Figure 4D). In line with the TRAP activity measurements, highest cathepsin K activity was found for experimental run 1, with a statistically significant difference with both positive and negative control (Figure 4F). From SEM images, osteoclast-like cells were observed in run 1 - 8 (Figure 4G). No apparent osteoclast-like cells (*i*.*e*., >10 μm in diameter and a ruffled boarder) were found in the negative control (run 9). In run 1 and 7, relatively large osteoclast-like cells were observed and in run 1 these cells were found in groups. Osteoclast-like cells in other experimental runs were generally smaller than osteoclast-like cells found in run 1 and 7 (Figure 4G).

**Figure 4.**
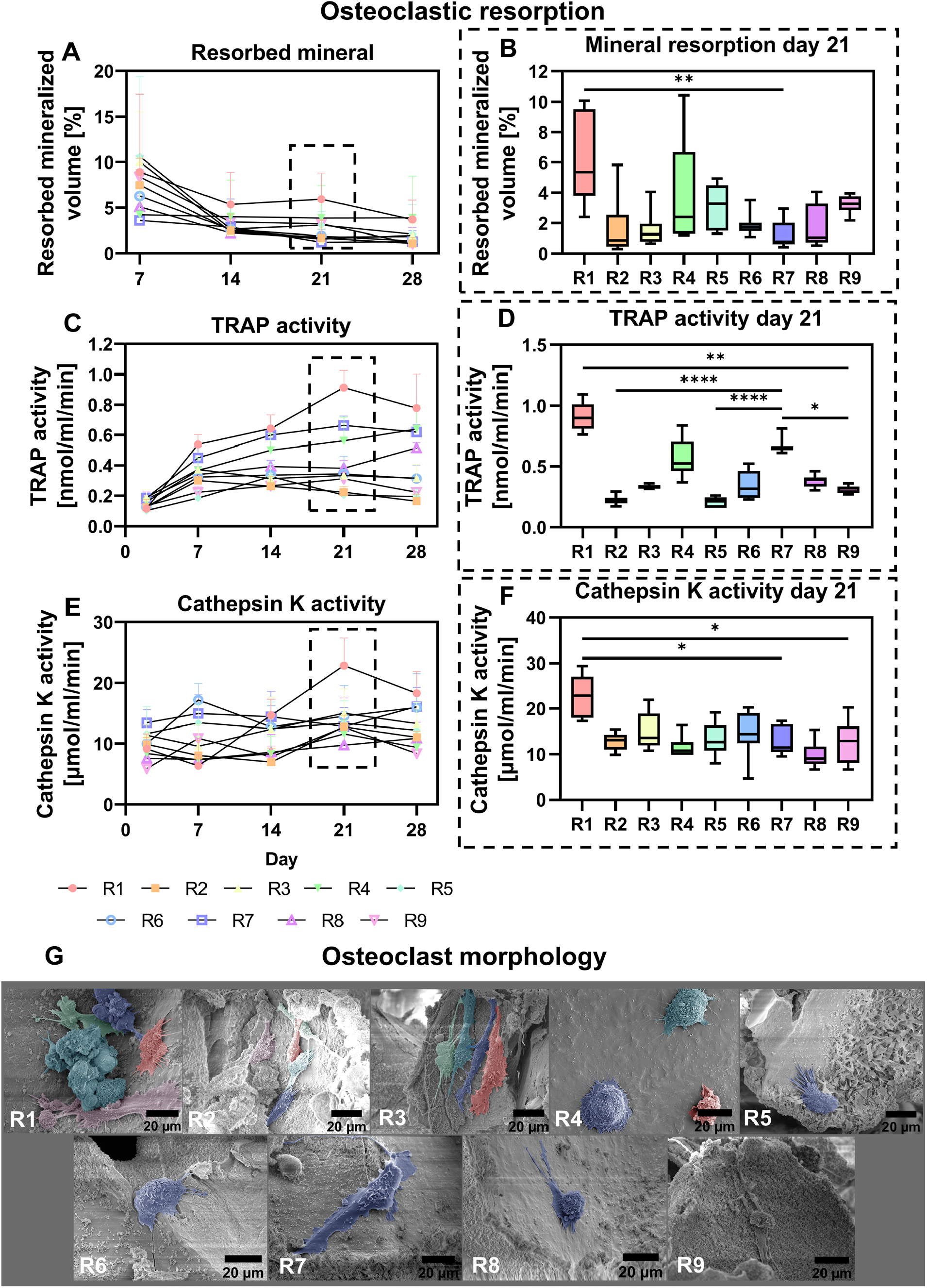
Osteoclastic resorption of experimental runs. (**A**) μCT based resorbed mineral of day 2-7, 7-14, 14-21, and 21-28. (**B**) Day 14-21 resorbed mineral, *p*<0.05 (Kruskal-Wallis and Dunn’s post hoc tests). (**C**) TRAP activity in the medium on day 2, 7, 14, 21, and 28. (**D**) Day 21 TRAP activity measurements, *p*<0.05 (Kruskal-Wallis and Dunn’s post hoc tests). (**E**) Cathepsin K activity in the medium on day 2, 7, 14, 21, and 28. (**F**) Day 21 cathepsin K activity measurements, *p*<0.05 (Kruskal-Wallis and Dunn’s post hoc tests). (**G**) Visualization of osteoclast-like cells on day 28 with SEM. (\**p*<0.05, \*\**p*<0.01, \*\*\*\**p*<0.0001). Abbreviations: tartrate-resistant acid phosphatase (TRAP), run (R), scanning election microscopy (SEM).

Factor main effect plots indicated the influence of the culture variables and their corresponding levels on mineral resorption, TRAP activity and cathepsin K activity (Figure 5). No significant contribution to mineral resorption, TRAP activity or cathepsin K activity was found from one of the factors. From the main effect plots and normal effect plots, mechanical loading tended to positively influence mineral resorption and TRAP activity, while the addition of high concentrations of osteogenic supplements tended to negatively influence mineral resorption (Figure 5 and Figure S2). Overall, factor main effect plots showed a similar trend for almost each resorption outcome measure (Figure 5). Only TRAP activity was positively influenced by a high concentration of hPL while resorbed mineral and cathepsin K were negatively influenced by a high concentration of hPL (Figure 5). This might be explained by the presence of TRAP in hPL [12].

**Figure 5.**
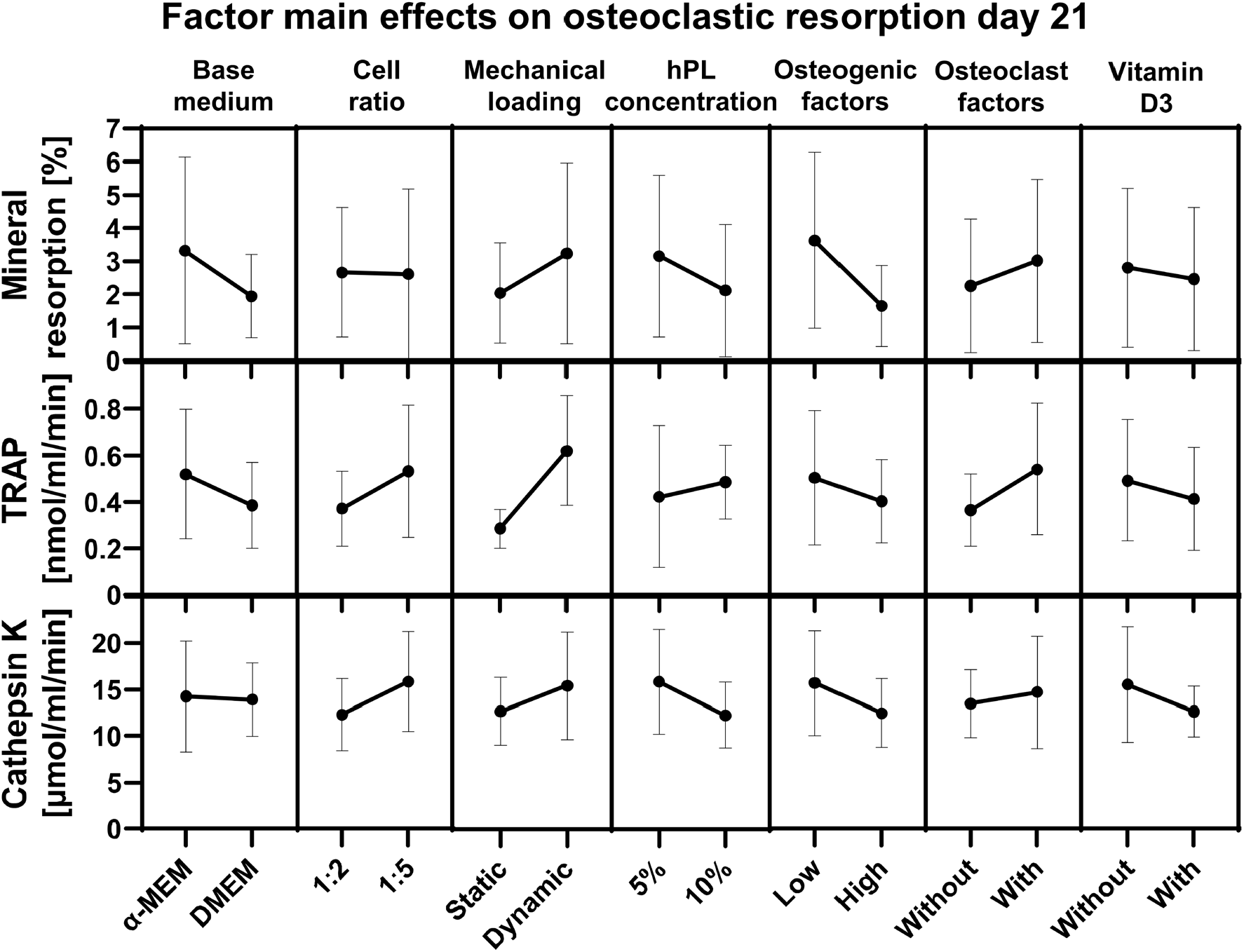
Factor main effects and standard deviations on day 21 osteoclastic resorption outcome measures. Abbreviations: tartrate-resistant acid phosphatase (TRAP), human platelet lysate (hPL).

### 3.4 Osteoblastic formation

When evaluating formed mineralized volume over time, the positive control (run 7) tended to have consistently high mineral formation while in the negative control (run 9), a relatively low formed mineralized volume was observed (Figure 6A). On day 21, only for run 1 a statistically significant higher formed mineralized volume was observed when compared to the negative control (Figure 6B). Similar trends were observed for collagen type 1 formation, measured by PICP release in the cell supernatant. Where most mineral formation was observed in run 1 and 7, also highest collagen type 1 formation was observed in these conditions (Figure 6C+D). In the negative control, limited collagen type 1 formation was observed. ALP activity measurements on the cell lysate of day 28 revealed highest ALP activity in the positive control and lowest in the negative control (Figure 6E). For the positive control, a statistically significant higher ALP activity was observed when compared to the ALP activity of run 5 and the negative control. Visualization of the constructs with confocal microscopy demonstrated collagen formation in run 1, 3, 4, 5 and 6 (Figure 6F). Remarkably, although relatively high PICP was observed in the positive control, almost no collagen was observed in the microscopy samples (Figure 6F). In all conditions, hydroxyapatite was mainly observed on the SF scaffold trabeculae rather than in the by the cells produced extracellular matrix (Figure 6F). In run 8 and 9, limited numbers of cells were observed (Figure 6F).

**Figure 6.**
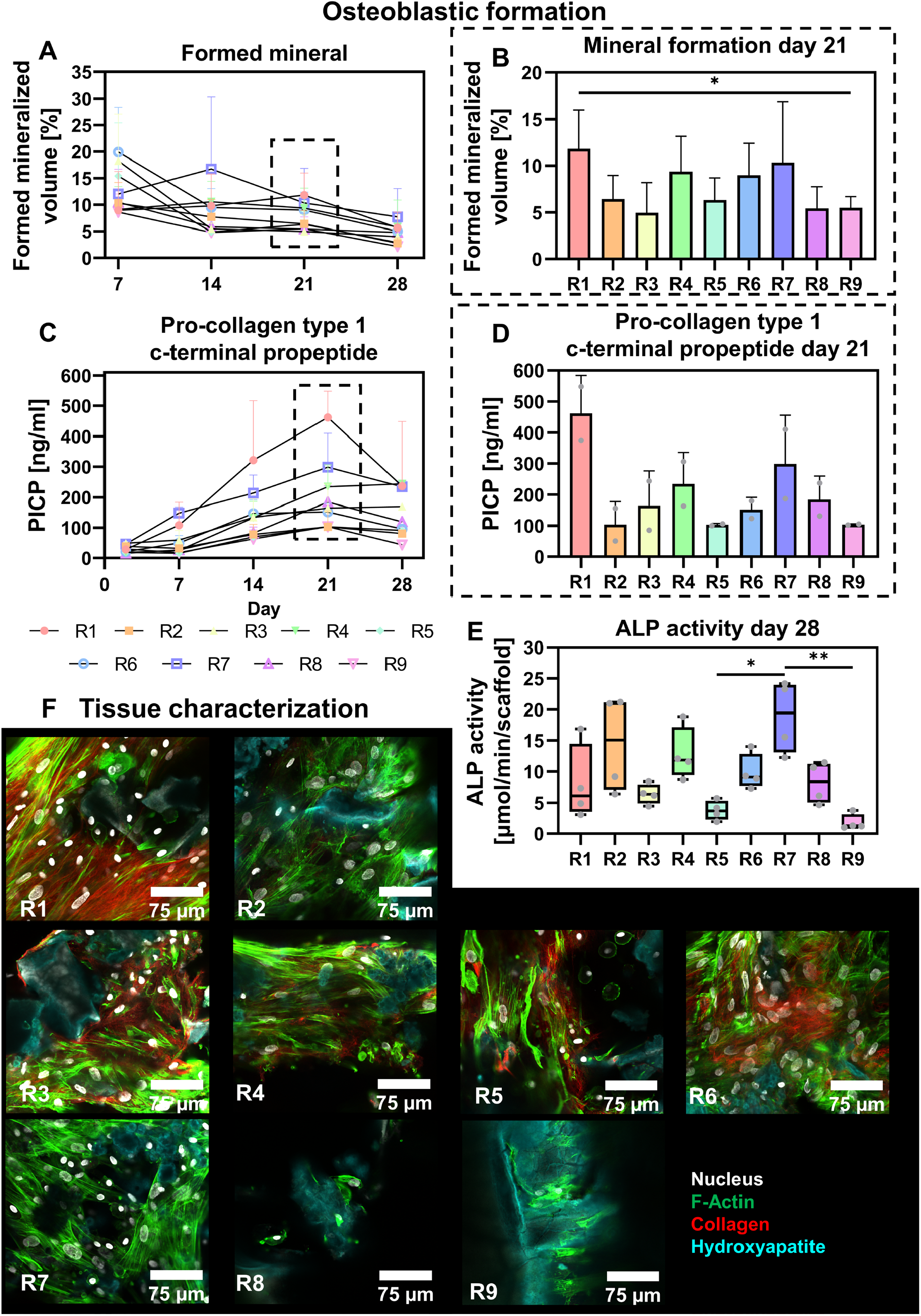
Osteoblastic formation of experimental runs. (**A**) μCT based formed mineral of day 2-7, 7-14, 14-21, and 21-28. (**B**) Day 14-21 formed mineral, *p*<0.05 (One-way ANOVA and Holm-Šídák’s post hoc tests). (**C**) PICP release in medium from day 2, 7, 14, 21, and 28. (**D**) Day 21 PICP release measurements. (**E**) ALP activity in cell lysates on day 28, *p*<0.05 (Kruskal-Wallis and Dunn’s post hoc tests). (**F**) Visualization of bone-like tissue in cultured constructs on day 28 with confocal microscopy, stained for collagen (red), F-Actin (green), hydroxyapatite (cyan), and the nucleus (gray). (\**p*<0.05, \*\**p*<0.01). Abbreviations: pro-collagen 1 c-terminal propeptide (PICP), alkaline phosphatase (ALP), run (R).

Factor main effect plots indicated the influence of the culture variables and their corresponding levels on mineral formation, collagen type 1 formation measured by PICP, and ALP activity (Figure 7A). No significant contribution to mineral formation, collagen type 1 formation, and ALP activity was found from one of the factors. From the main effect plots and normal effect plots, mechanical loading tended to positively influence mineral formation and collagen type 1 formation (Figure 7A and Figure S2). Other factors did not seem to influence mineral formation (Figure 7A and S2). ALP activity tended to be positively influenced by high concentrations of osteogenic supplements (Figure 7A and S2). As mineral formation was relatively high in groups where mineral resorption was elevated in the absence of high stimulation with osteogenic factors, we investigated the coupling between resorption and formation markers (Figure 7B+C). Interestingly, a strong positive correlation (r = 0.81, *p* < 0.0001) was observed between the osteoclastic resorption marker TRAP and the collagen formation marker PICP (Figure 7B). In line with these results, main effect plots and normal effect plots for TRAP activity and PICP concentration followed a similar pattern (Figure 5, 7A and S2), which could suggest an influence of osteoclastic differentiation and/or TRAP activity on osteoblastic collagen formation. When investigating the coupling between mineral resorption and formation, a moderate positive correlation (r = 0.59, *p* < 0.0001) was found (Figure 7C). When splitting the data into high and low stimulation with osteogenic factors, a weak positive (r = 0.38, *p* < 0.05) correlation between mineral resorption and formation was found in highly stimulated scaffolds whereas a strong positive correlation (r = 0.80, *p* < 0.0001) was found when low stimulation was applied (Figure 7C). This indicates that resorption - formation coupling can be disturbed by high stimulation with osteogenic factors.

**Figure 7.**
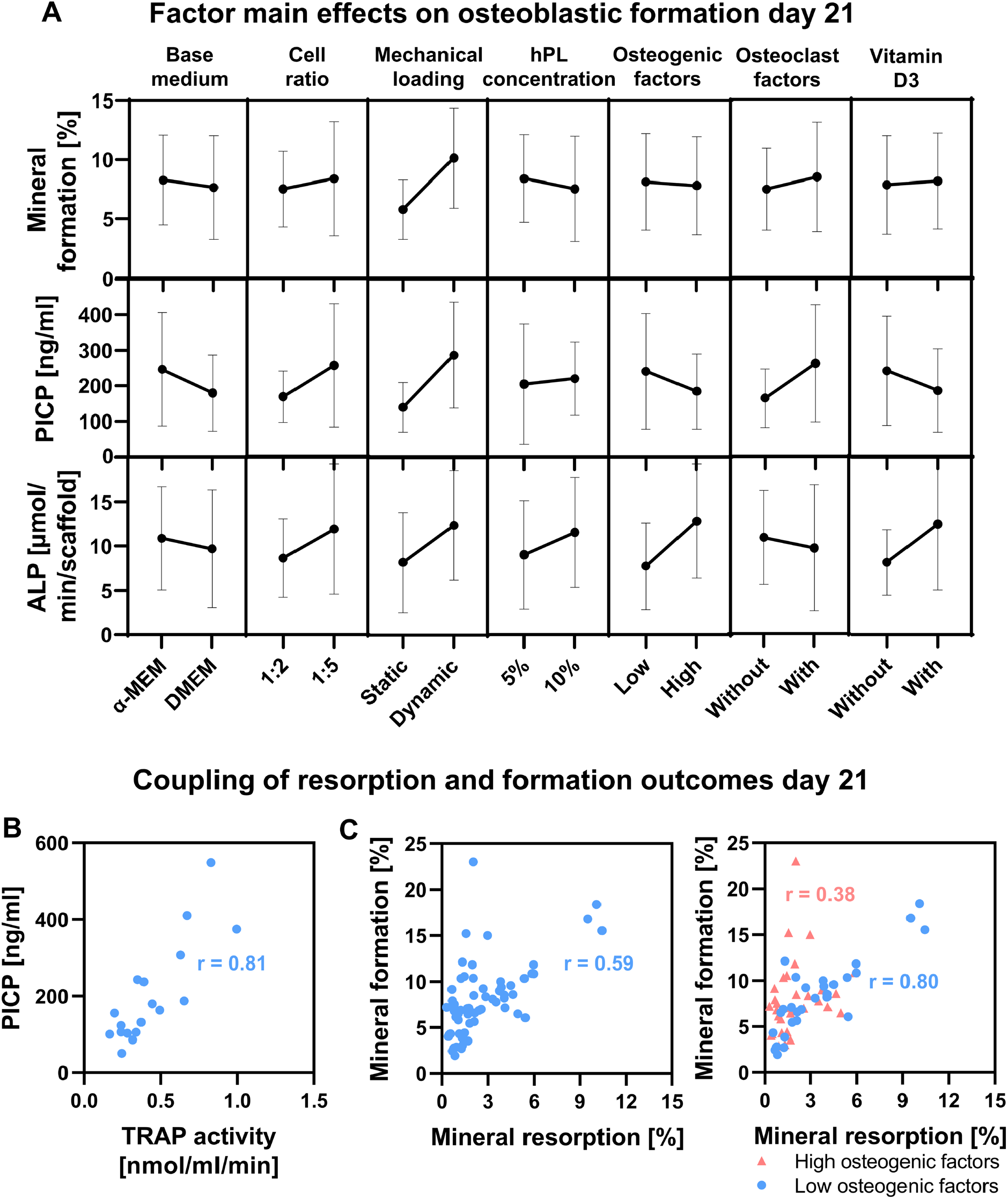
(**A**) Factor main effects and standard deviations on day 21 osteoblastic formation outcome measures. Correlation/coupling of resorption and formation outcomes for (**B**) organic matrix resorption and formation (TRAP and PICP), and (**C**) inorganic matrix resorption and formation, in the presence and absence of high stimulation with osteogenic differentiation factors. Abbreviations: human platelet lysate (hPL), tartrate-resistant acid phosphatase (TRAP), pro-collagen 1 c-terminal propeptide (PICP).

## 4. Discussion

Human *in vitro* bone remodeling models, using osteoclast-osteoblast co-cultures, could facilitate the investigation of human healthy (*i*.*e*., balanced) and pathological (*i*.*e*., unbalanced) bone remodeling while addressing the principle of 3Rs for animal experiments [4,5]. Although current *in vitro* osteoclast-osteoblast co-cultures have improved our understanding of bone remodeling, they lack standardization of culture methods and outcome measurements, hampering reproducibility and translatability to *in vivo* animal models and *in vivo* human data. In this regard, *in vitro* bone remodeling models could benefit from a thorough evaluation of the impact of culture variables on functional and translatable outcome measures, with the aim to reach ‘healthy’ balanced osteoclast and osteoblast activity. Using a resolution III fractional factorial design, we identified the main effects of commonly used culture variables at high and low stimulation on mineral resorption, mineral formation and a multitude of bone turnover biomarkers in our *in vitro* human bone remodeling model.

We first evaluated the influence of culture variables on cell viability. None of the factors had a significant influence on cell metabolic activity or cell death. The absence of significant factors indicates that a combination of multiple factors contributed to the found differences between the experimental runs. Metabolic activity tended to decrease with high stimulation of all factors other than cell ratio. As the initial seeding density of hBMSCs was the same for the different experimental runs, a higher cell ratio led to an increase in hMC density and thereby likely positively influencing cell metabolic activity. The negative influence on metabolic activity of the other culture variables with high stimulation might be explained by differences in energy metabolism of undifferentiated and differentiated progenitor cells. In this study, metabolic activity was measured by the reduction of resazurin to fluorescent resorufin by aerobic respiration. In contrast to our metabolic activity results indicating less metabolic activity upon cell differentiation, osteogenic differentiation of hMSCs has shown to increase the portion of aerobic respiration to the cells’ energy metabolism [35]. Moreover, osteoclast differentiation is associated with increased mitochondrial biosynthesis and oxygen consumption rate, likely enhancing aerobic respiration [36,37]. Thus, osteogenic and osteoclastic differentiation were expected to increase metabolic activity. Although high stimulation did not always lead to improved cell differentiation as measured with enzymatic activity markers (*i*.*e*., ALP, TRAP and Cathepsin K), one hypothesis for this contradiction might be that limited exogenous factor stimulation enhances endogenous factor production. To confirm this hypothesis, further cell secretome quantification using for example multiplex ELISA is required. If cells with limited stimulation are indeed producing more endogenous factors, limited stimulation might be essential to create self-regulating models.

In our effort to mimic healthy (balanced) remodeling *in vitro*, the influences of culture variables on bone turnover parameters were evaluated. Although the negative control showed most balanced resorption and formation, only limited remodeling could be detected, while the ability to capture remodeling is imperative for *in vitro* remodeling models. On the other hand, the positive control showed least balanced resorption and formation, with a relatively high net formation and low volume of quiescent mineral. As such, both low and high stimulation are non-optimal to mimic bone homeostasis *in vitro*. With the least quiescent mineral, experimental run 1 (*α*-MEM, 1:5, dynamic, 5% hPL, osteoclast factors) stood out. When evaluating the resorption dynamics of this experimental run, most mineral resorption and highest activity of matrix degradation enzymes TRAP and Cathepsin K were measured, and typical osteoclast-like cells were identified. Therefore, run 1 is considered most optimal for *in vitro* bone resorption. Remarkably, in terms of osteoblastic formation, run 1 also showed most mineral and collagen type 1 formation. Only for ALP activity in the cell lysates, run 1 showed levels in between the positive and negative control. This appears contradictory but could well be due to downregulation of ALP activity upon fast osteoblast maturation [38].

Thus far, the use of exogenous osteoclast differentiation factors (*i*.*e*., RANKL and M-CSF) in osteoclast-osteoblast co-cultures is generally considered crucial for the development of functional osteoclasts [4], even though exogenous application of these factors could overrule the natural RANKL/osteoprotegerin (OPG) ratio as important regulator in healthy and pathological bone remodeling [4,39]. Strikingly, when we replaced osteoclastogenic and osteogenic differentiation factors by 1,25-dihydroxyvitamin D3 (run 4; *α*-MEM, 1:2, dynamic, 10% hPL, 1,25-dihydroxyvitamin D3), we observed relatively high osteoclast and osteoblast activity. Even in the absence of differentiation factors, relatively high levels of mineral resorption, TRAP activity, mineral formation, collagen type 1 production, and ALP activity were found. When no differentiation factors were used in human PBMC-osteoblast co-cultures, no osteoclastic differentiation of PBMCs was observed [40]. Other researchers found that when human PBMCs and hBMSCs were co-cultured on osteoblast derived matrix in the absence of osteogenic and osteoclastic differentiation factors, resorption was comparable to PBMC mono-cultures treated with M-CSF and RANKL [41]. Moreover, before the discovery of RANKL and the ability to clone this factor, stromal cells and osteoblasts were used as a tool for osteoclastic differentiation *in vitro* [42]. We therefore believe that osteoclastic differentiation in osteoclast-osteoblast co-culture in the absence of exogenous RANKL is possible under the correct circumstances. With the use of 1,25-dihydroxyvitamin D3 in experimental run 4, RANKL expression by hMSCs/osteoblasts and subsequent osteoclastic differentiation might have been stimulated as earlier demonstrated [42,43]. It would be interesting to quantify M-CSF, RANKL and OPG produced in the experimental runs to investigate the influence of 1,25-dihydroxyvitamin D3 on the osteoclastic differentiation potential in our co-cultures, with the ambition to circumvent the use of exogenous osteoclastic differentiation factors in future.

The enhanced resorption and formation in experimental run 1 raised the expectation that resorption and formation were coupled in our *in vitro* model. Indeed, resorption and formation were correlated for both mineral resorption/formation and organic matrix resorption/formation. By adding an exogeneous phosphate source to the model (*β*-glycerophosphate), mineral resorption – formation coupling was disturbed. Most likely, osteoclasts released sufficient calcium phosphate from the mineralized scaffold for subsequent formation. *In vivo*, coupling includes communication through secreted and cell-bound factors, topographical ques, and the release of growth factors from the bone matrix, with the main goal to replace the resorbed bone volume by an equal volume of new bone [44–46]. In addition, osteocytes as well as osteoclast and osteoblast progenitors contribute to this coupling, which likely changes their contribution during their differentiation towards mature osteoclasts and osteoblasts [47,48]. As such, coupling is a highly complex process, and it is expected that not all coupling aspects are present in our model. More specifically, the release of growth factors from the matrix/scaffold and the contribution of osteocytes are likely lacking. To enable coupling through growth factor release, *in vitro* remodeling models could be developed on decellularized bone tissue [49,50]. To additionally involve osteocytes into the bone remodeling process *in vitro*, a long-term pre-culture in which osteoprogenitors differentiate into osteocytes while they develop their mineralized niche [51–53], or the use of cell-lines might be required, due to challenges with primary osteocyte isolation and subsequent culture [54]. To combine the presence of a growth-factor containing bone matrix and osteocytes, osteocytes could also be cultured in their native niche using human trabecular bone specimens [54]. In this study we found quantitative coupling between resorption and formation at the tissue level. To further validate coupling in our *in vitro* model, it would be interesting to study coupling qualitatively at the level of the individual resorption pits to see whether formation takes place on previously resorbed surfaces like in the model of A. Hikita *et al*, (2015) [55]. Nevertheless, quantitative coupling was observed along all conditions, indicating some endogenous regulation in all conditions which can be enhanced by external stimuli.

The application of mechanical loading tended to be the most influential factor on both resorption and formation outcomes. While mechanical loading in terms of fluid flow induced shear stress has, in line with our results, been shown to stimulate mineralization and collagen formation in similar settings [21,52], its clear effect on resorption outcomes was unexpected. *In vivo*, bone remodeling and adaptation is regulated by osteocytes under influence of interstitial fluid flow through the lacuno-canalicular network [56]. Osteocytes that sense mechanical loading could inhibit osteoclastic differentiation both directly and indirectly [57]. The direct influence of mechanical loading on osteoclast differentiation is relatively unknown with *in vitro* both positive [58,59] and negative [60,61] influences reported in literature. As mentioned above, the contribution of osteocytes and thereby their inhibitory influence on resorption under influence of mechanical loading is likely lacking. Another explanation for the enhanced resorption under influence of mechanical loading could be the improved mass transport when fluid flow was applied. It would therefore be interesting to study the interaction between osteoclastic differentiation factors and mechanical loading within our model, to check whether the likely improved distribution of osteoclastic differentiation factors indeed leads to increased osteoclastic differentiation.

One limitation of the current set-up is that four scaffolds are placed within one bioreactor. Medium analyses therefore only show the average per bioreactor containing two donor combinations. Another limitation of the current study is the resolution of the fractional factorial design. With the use of a resolution III design, only an influence of factor main effects could be provided. Moreover, some of these main effects are confounded with interaction effects, which complicates outcome interpretation. In our evaluation, we did not find significant contributions of specific factors to cell viability and bone turnover outcomes. Nevertheless, clear differences between experimental runs were observed. This suggests that a combination of multiple factors contributed to the found differences between experimental runs. Additionally, by both using three different hMC and hBMSC donors, and three different methods for each resorption and formation, we here present a robust evaluation of the influence of culture variables on *in vitro* bone remodeling.

## 5. Conclusion

With the aim to mimic healthy balanced bone remodeling *in vitro*, we have identified the impact of commonly used culture variables on translatable bone turnover parameters in a human bone remodeling model. We herewith present a robust *in vitro* bone remodeling model, that was able to capture physiological quantitative resorption – formation coupling along all conditions, which could be enhanced by external stimuli. As such, *in vitro* remodeling (*i*.*e*., resorption and formation) was enhanced by the application of mechanical loading. Moreover, high stimulation with osteogenic differentiation factors disturbed mineral resorption – formation coupling. Especially culture conditions of run 1 (*α*-MEM, 1:5, dynamic, 5% hPL, osteoclast factors) and run 4 (*α*-MEM, 1:2, dynamic, 10% hPL, 1,25-dihydroxyvitamin D3) showed promising results, where run 1 conditions could be used as high bone turnover system and run 4 as self-regulating system as the addition of osteoclastic and osteogenic differentiation factors was not required for remodeling. The results generated with our *in vitro* model allow for better translation between *in vitro* studies and towards *in vivo* studies, for improved preclinical bone remodeling drug development.

## 7. Author Contributions

BdW, LC, KI and SH contributed to conception and design of the study. BdW, EC and SH contributed to the methodology. BdW performed the experiments. BdW and AW analysed the experimental results. BdW wrote the original draft of the manuscript. BdW, KI and SH were involved in supervision. All authors contributed to manuscript revision and approved the submitted version. BdW and SH acquired funding for this research.

