## Supplementary Material for "The impact of culture variables on 3D human *in vitro* bone remodeling; a design of experiments approach"

Each experimental run in the fractional factorial design had a unique medium composition (Table S1 and S2).

**Table S1.** Medium compositions for different experimental runs.

| | base medium | hPL | anti-anti | HEPES | L-glut | ascorbic acid | β-GP | dex | RANKL * | M-CSF | vit D3 |
|---|---|---|---|---|---|---|---|---|---|---|---|
| R1 | 94% α-MEM | 5% | 1% | 25 mM | | 50 μg/ml | | 10 nM | 50 ng/ml | 50 ng/ml | |
| R2 | 94% α-MEM | 5% | 1% | 25 mM | | 50 μg/ml | 10 mM | 100 nM | | | 10 nM |
| R3 | 89% DMEM | 10% | 1% | 25 mM | 4mM | 50 μg/ml | | 10 nM | | | |
| R4 | 89% α-MEM | 10% | 1% | 25 mM | | 50 μg/ml | | 10 nM | | | 10 nM |
| R5 | 94% DMEM | 5% | 1% | 25 mM | 4mM | 50 μg/ml | | 10 nM | 50 ng/ml | 50 ng/ml | 10 nM |
| R6 | 94% DMEM | 5% | 1% | 25 mM | 4mM | 50 μg/ml | 10 mM | 100 nM | | | |
| R7 | 89% DMEM | 10% | 1% | 25 mM | 4mM | 50 μg/ml | 10 mM | 100 nM | 50 ng/ml | 50 ng/ml | 10 nM |
| R8 | 89% α-MEM | 10% | 1% | 25 mM | | 50 μg/ml | 10 mM | 100 nM | 50 ng/ml | 50 ng/ml | |
| R9 | 94% α-MEM | 5% | 1% | 25 mM | | 50 μg/ml | | 10 nM | | | |

Abbreviations: run (R), human platelet lysate (hPL), antibiotic antimycotic (anti-anti), L-glutamine (L-glut), β-glycerophosphate (β-GP), dexamethasone (dex), macrophage colony-stimulating factor (M-CSF), receptor activator of nuclear factor κB ligand (RANKL), 1,25-dihydroxyvitamin D3 (vit D3).
*applied from day 2 in culture

**Table S2.** Medium components and suppliers.

| | |
|---|---|
| α-MEM | 41061, Thermo Fisher Scientific, Breda, The Netherlands |
| DMEM | 11880, Thermo Fisher Scientific |
| hPL | PE20612, PL BioScience, Aachen, Germany |
| Anti-anti | 15240, Thermo Fisher Scientific |
| HEPES | 15630, Thermo Fisher Scientific |
| L-glutamin | X0550-100, Biowest, Nuaillé, France |
| Ascorbic acid-2-phosphate | A8960, Sigma Aldrich, Zwijndrecht, The Netherlands |
| β-glycerophosphate | G9422, Sigma-Aldrich |
| Dexamethasone | D4902, Sigma-Aldrich |
| RANKL | 310-01, PeproTech, London, UK |
| M-CSF | 300-25, PeproTech |
| 1,25-dihydroxyvitamin D3 | D1530, Sigma-Aldrich |

Abbreviations: human platelet lysate (hPL), antibiotic antimycotic (anti-anti), macrophage colony-stimulating factor (M-CSF), receptor activator of nuclear factor κB ligand (RANKL).

Normal effect plots were generated to visualize the effect size and direction of each factor on cell viability (Figure S1) and bone turnover outcomes (Figure S2).

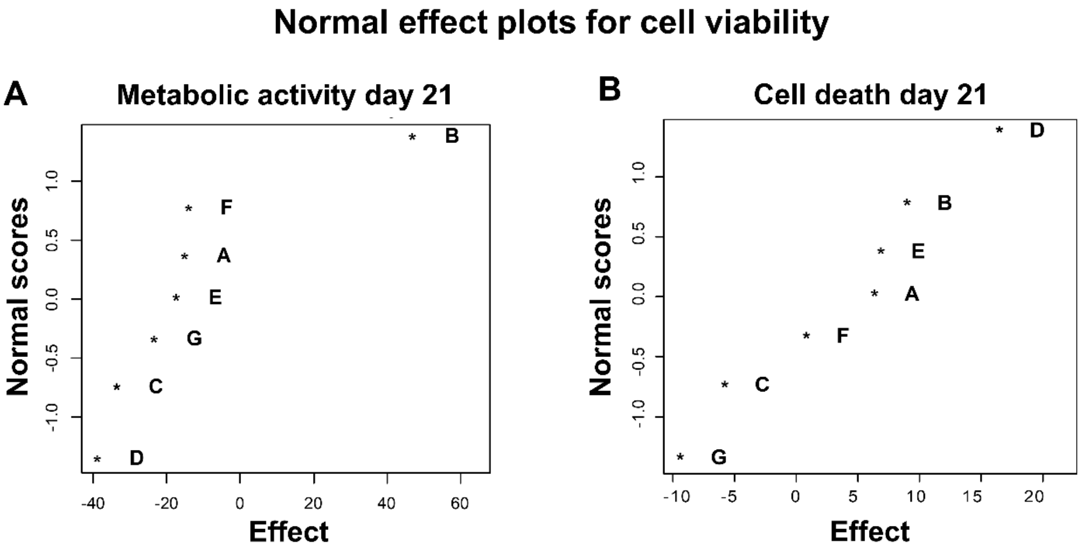


**Figure S1.** Normal effect plots for cell viability outcomes. (**A**) Normal effect plot for day 21 metabolic activity, indicating a non-significant positive effect of a high cell ratio on metabolic activity. (**B**) Normal effect plot for day 21 cell death. A = base medium, B = cell ratio, C = mechanical loading, D = human platelet lysate concentration, E = osteogenic factors, F = osteoclast factors, G = 1,25-dihydroxyvitamin D3

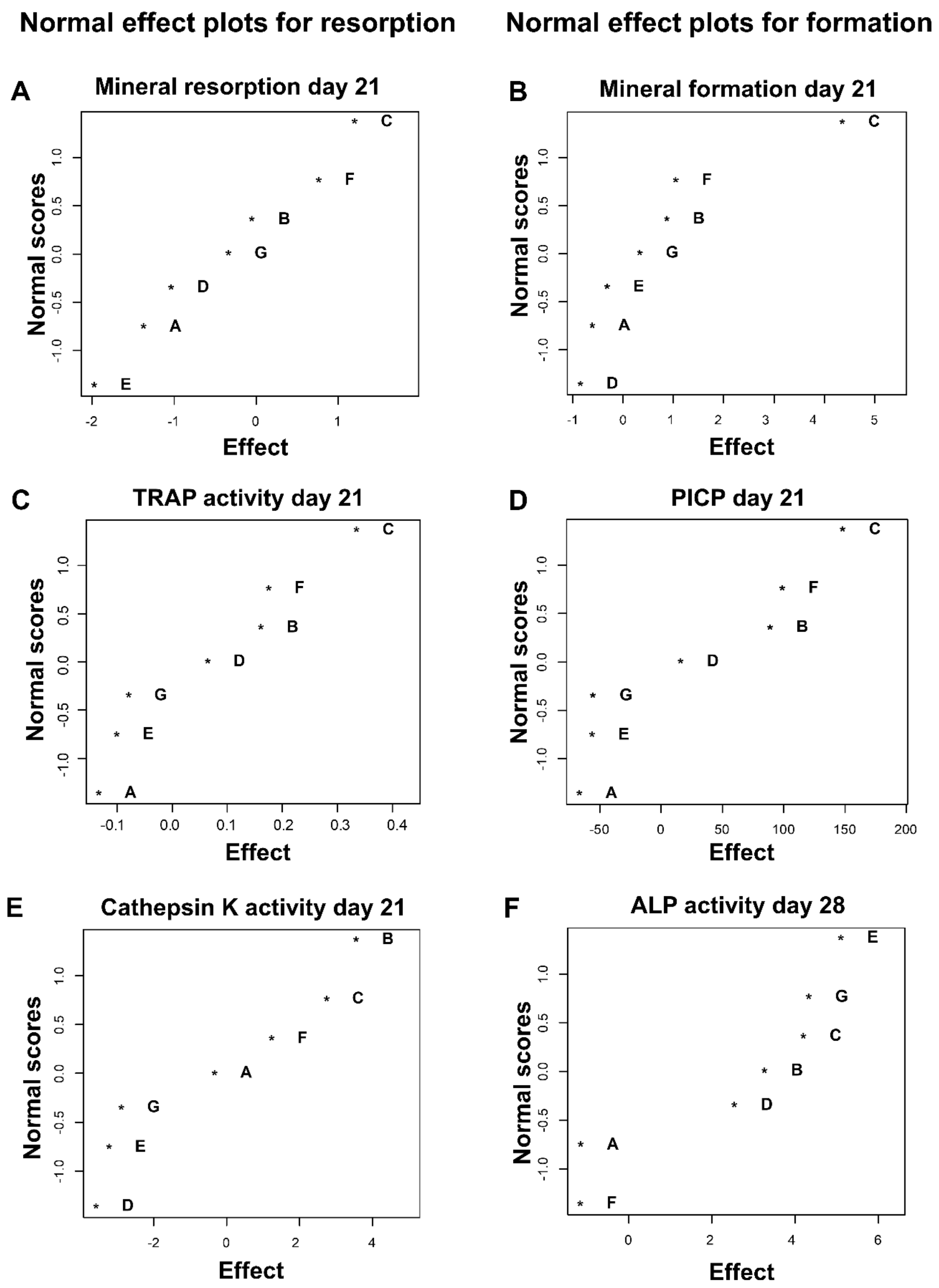


**Figure S2.** Normal effect plots for bone turnover outcomes. (**A**) Normal effect plot for day 21 mineral resorption. (**B**) Normal effect plot for day 21 mineral formation, indicating a non-significant positive effect of high stimulation with mechanical loading on mineral formation. (**C**) Normal effect plot of day 21 TRAP activity. (**D**) Normal effect plot of day 21 PICP. (**E**) Normal effect plot of day 21 Cathepsin K activity. (**F**) Normal effect plot of day 28 ALP activity. A = base medium, B = cell ratio, C = mechanical loading, D =

human platelet lysate concentration, E = osteogenic factors, F = osteoclast factors, G = 1,25-dihydroxyvitamin D3. Abbreviation: tartrate-resistant acid phosphatase (TRAP), pro-collagen 1 c-terminal propeptide (PICP), alkaline phosphatase (ALP).
